# *ChoruMM*: a versatile multi-components mixed model for bacterial-GWAS

**DOI:** 10.1101/2023.03.28.534531

**Authors:** Arthur Frouin, Fabien Laporte, Lukas Hafner, Mylene Maury, Zachary R. McCaw, Hanna Julienne, Léo Henches, Rayan Chikhi, Marc Lecuit, Hugues Aschard

## Abstract

Genome-wide Association Studies (GWAS) have been central to studying the genetics of complex human outcomes, and there is now tremendous interest in implementing GWAS-like approaches to study pathogenic bacteria. A variety of methods have been proposed to address the complex linkage structure of bacterial genomes, however, some questions remain about to optimize the genetic modelling of bacteria to decipher causal variations from correlated ones. Here we examined the genetic structure underlying whole-genome sequencing data from 3,824 *Listeria monocytogenes* strains, and demonstrate that the standard human genetics model, commonly assumed by existing bacterial GWAS methods, is inadequate for studying such highly structured organisms. We leverage these results to develop *ChoruMM*, a robust and powerful approach that consists of a multi-component linear mixed model, where components are inferred from a hierarchical clustering of the bacteria genetic relatedness matrix. Our *ChoruMM* approach also includes post-processing and visualization tools that address the pervasive long-range correlation observed in bacteria genome and allow to assess the type I error rate calibration.

## Introduction

The Genome-wide Association Study (GWAS) is an experimental design that has been developed to detect associations between genetic variants, usually single-nucleotide polymorphisms (SNPs), and traits in samples from populations. GWAS has become the primary tool to study highly polygenic phenotypes in humans^1^ but also other mammals^2^ and plants^3^. Its widespread use has arisen thanks to its simplicity and robustness, the intense effort for developing efficient genotyping technologies, but also to the underlying rationale: the systematic search for variants achieving a stringent significance level has proven critical to ensuring high replicability of results, an essential feature of scientific and public health research. With the continuous reduction in sequencing costs, there is growing interest in applying GWAS-like approaches to the genomes of pathogenic bacteria in order to advance our understanding of infectious disease risk and identify genetic variants driving bacterial virulence^4-7^. This interest come along extended discussion on how to adapt the GWAS methodology to bacteria. Early studies showed that the standard approach used in human genetics, is in theory applicable to core genome mutations in bacteria (e.g.^8-10^), but also outlined some limitations. Indeed, bacterial genomes vary extensively in terms of both gene content and gene sequences. Because of limited recombination, bacterial genomes also harbor large linked haplotype blocks inducing long-range linkage disequilibrium (LD). Moreover, many bacterial genomes do not possess large core genomes, and strains’ sequences are not always aligned, hindering the use of traditional single nucleotide polymorphisms (SNP)-based methods^4^.

Important works have explored strategies to adapt GWAS for bacterial genetics and paved the way for future developments^6,11^. Putting aside approaches using *ad hoc* adjustment for population structure (*e*.*g*. phylogenic information as in *TreeWAS*^*12*^ and *phyOverlap*^*13*^) and permutation-based approaches (e.g. *Scoary*^*14*^), approaches using a formal statistical framework have focused on two critical aspects: 1) how to express the diversity of genetic variations within a single unified framework, and 2) how to account for population structure within the inference model. For genetic variation, the community has partly converged toward using DNA words of length *k*^15^, referred as *k-mers*, as a generalized alternative to SNPs, thanks to their ability to accommodate complex genomic structure, including rearrangements, insertions, and deletions^7^. Multiple tools have been developed to perform *k-mer* based GWAS (e.g. *bugwas*^*16*^, *HAWK*^*17,18*^). *k-mers* have the primary advantage of not requiring a reference genome, nor genome assembly. On the other hand, they suffer strong redundancy and can be difficult to interpret. To solve this issue, Jaillard et al^19^ proposed using compacted De Bruijn Graphs (cDBGs) to connect overlapping *k*-mers to form unitigs, including SNPs, insertion/deletion, and presence/absence of genes or any long sequence. Regarding the genetic structure, two primary approaches have been considered (sometimes conjointly^20^), using the first principal components (PCs) from a spectral decomposition of the genetic relationship matrix^18^, and using a genetic relatedness matrix introduced via a linear mixed model (LMM). Thanks to its versatility and successful application in humans^21^, the LMM approach is becoming increasingly adopted for bacterial GWAS and has been included in several packages including *bugwas*^*16*^, *pySEER*^*22*^ and *DBGWAS*^*19*^. Nevertheless, questions remain regarding the differences between bacterial and human genome pertaining to the GWAS approach, and about the impact of model parameterization on our ability to fully address the complex structure underlying bacterial genetics.

Here we show that some characteristics of the bacterial genomes are not handled by the standard GWAS approach. We discuss how these gaps impact reliability, applicability and interpretability, three factors that made the human GWAS efficient and popular. Building over these results and the state-of-the-art, we develop *ChoruMM* a robust, powerful and versatile multi-component linear mixed model (MLMM) that improves the modelling of complex bacteria structure. We incorporated *ChoruMM* into an end-to-end user-friendly tool including our MLMM and multiple *ad hoc* tools for pre-processing unitig data and visualizing bacterial GWAS results.

## Results

### Overview

We characterized bacterial genetic structure and develop our method using real whole-genome sequencing dataset of *Listeria monocytogenes* strains. All analyses were conducted using unitigs inferred from the raw sequence data with compacted De Bruin Graphs (**Supplementary Notes** and **Fig. S1**). We first use those data to characterize the extend of linkage disequilibrium across variants, the distribution of relatedness across strains, and to examine the impact of this structure on heritability across Listeria lineages. We then present the *ChoruMM* approach, a multi-component linear mixed model (MLMM), that we benchmarked for both power and robustness using real unitigs and simulated phenotype data. Finally, we introduce multiple complementary innovations to assess the calibration of the type I error rate and to facilitate the visualization and interpretation of the GWAS results.

### From human to bacterial GWAS

The human and bacterial genomes differ in many ways. However, some global features typical of human GWAS analysis have been only seldom described in bacterial genomes. To guide the implementation of a GWAS approach, we conducted a series of analyses to characterize the extent of genetic structure in bacteria.

First, we quantified both the extend of relatedness between strains and the correlation between genetic variants. Strain relatedness is underscored by examining the cumulative distribution of eigenvalues derived from the unitig matrix in *Listeria*. As shown in **Figure 1a**, the top 10 principal components explain roughly 20% of the variance. In comparison, the top 10 principal components derived from genotypes in a standard human GWAS would typically explain less than 1% of the total genetic variation^18^, and thousands of components would be necessary to reach 20%. As for the correlation between genetic variants, this is illustrated using linkage disequilibrium (LD)-pruning, a common technique in human genetics which filters out variants with a squared-correlation (R^2^) above a given threshold. In a standard human GWAS dataset, pruning a set of (e.g.) 1M common variants from HapMap3 SNPs^19^ at an R^2^ of 0.05 results in roughly 50K independent SNPs. We plotted in **Figure 1b** the count of unitigs remaining after LD pruning with an R^2^ in [0.99, 0.05] for the Listeria cohort. Overall, less than 0.5% of the unitigs can be considered independent (R^2^ < 0.05). Even using a very lenient threshold of R^2^ < 0.5 results in the removal of over 90% of the unitigs.

**Figure 1.**
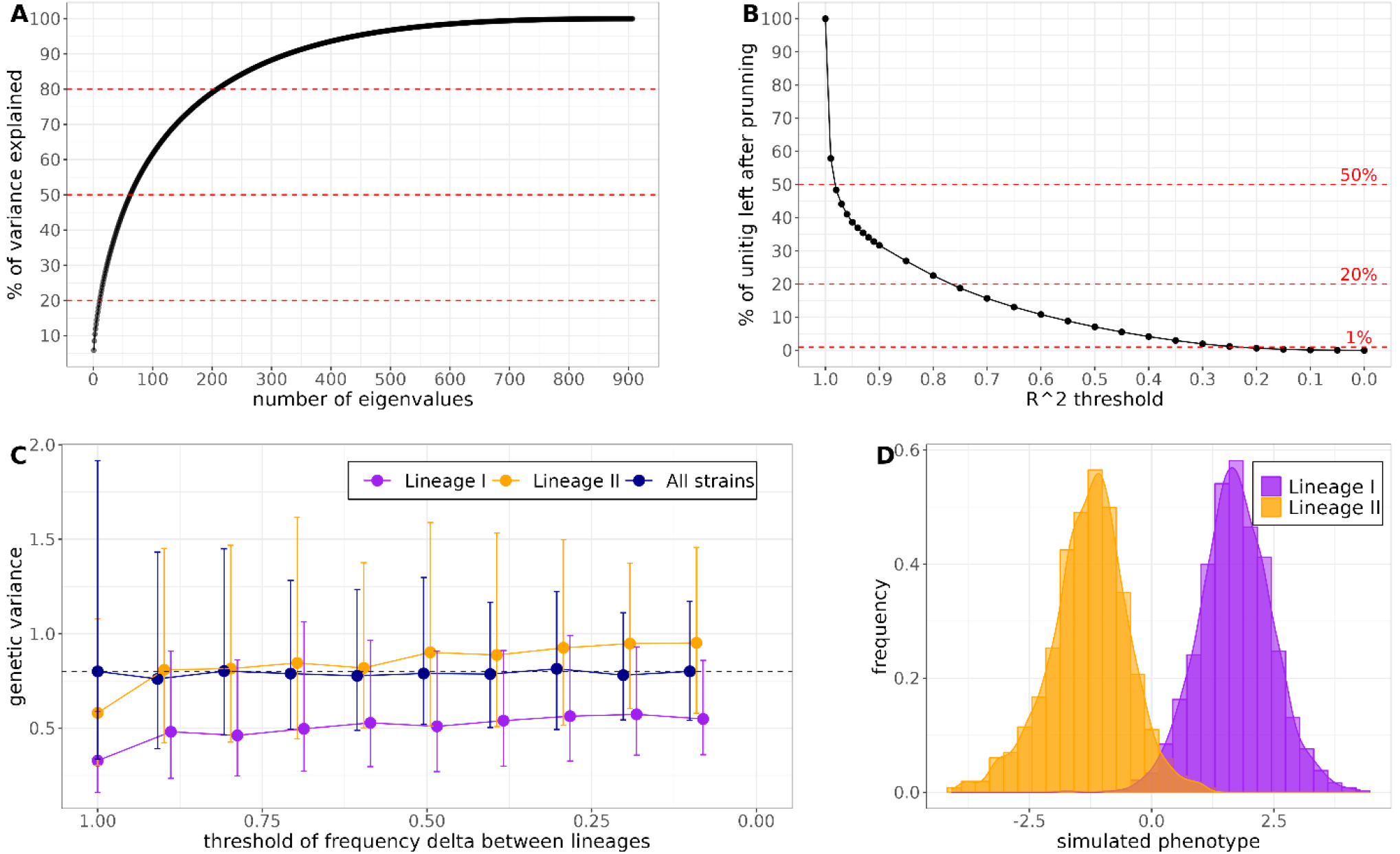
Realized Genetic component in highly structured bacterial data. Panel **A)** presents the cumulative percentage of variance explained by the eigenvalues from a unitig genetic relatedness matrix (GRM). The GRM of 907 strains was computed on a subset of pruned unitigs (R^2^ < 0.2). **B)** Proportion of unitigs remaining after pruning for various values of R^2^. The pruning was realized on a unitig matrix of 907 strains and 320,247 unitigs. **C)** Point estimate and 95% standard error of estimated genetic variance from simulated data for the whole sample (blue curve) and per lineage (yellow and purple) as a function of the difference in frequency between lineage I and lineage II. Phenotypes were simulated according to a linear model with a single genetic term, from a matrix of 3824 strains distributed into two lineages (1642 lineage I and 2181 lineage II).2000 unitigs were randomly selected as causal based on whether their difference in frequency between the lineages exceeded a defined threshold (x axis). The true heritability was fixed as 0.8. **D)** Example of the extreme difference in the realized genetic variance between lineages for a simulated phenotype pulled from panel C (threshold of frequency delta = 1).

Next, we assessed the variability in the realized genetic variance between the two main *Listeria* lineages (I and II)^23^ across simulated phenotypes. Those two lineages display substantial genetic variability, providing a typical scenario for the analysis of highly structured data. Phenotypes were generated assuming a polygenic model using a subset of 2,000 causal unitigs selected based on their differences in frequency between lineages I and II. In these simulations, the genetic variance is unbiased when measured in the full cohort, whatever the set of causal unitig selected. Conversely, the realized genetic variance within each lineage varies substantially, being orders of magnitude smaller than for the whole sample in many situations. Using only unitigs with high frequency in the total sample amplifies this pattern, while using only unitigs with high frequency in both lineages reduces it (**Fig. 1c** and **Fig. S2**). As illustrated in **Figure 1d**, these differences are mostly explained by the bi-modal genetic variance produced when jointly analyzing strains with large genetic differences. For example, a causal unitig with frequencies of 100% and 0% in lineages I and II, respectively, will produce variance in the whole sample. However, in lineage-specific analyses, this unitig carries no information and does not contribute to the outcome variance. This implies that such a genetic effect can only be detected when jointly analyzing the two lineages and that heritability, defined as the ratio of the genetic variance over the total outcome variance, can vary markedly depending on the samples used.

### The ChoruMM linear mixed model

To address the complex genetic structure across strains, we propose to model the genetic structure using a multi-component linear mixed model, instead of a single component as proposed in previous bacterial GWAS^16,19,22^. We first present testing procedures based on linear mixed model (LMM), starting from the widely used one-component LMM.

Let **y** ∈ *R*^*n*^ be a vector of a quantitative phenotype of interest, **Z** ∈ *M*_*n,p*_(*R*) a standardized incidence matrix of genetic variants (e.g., SNPs, genes, unitigs) with rows corresponding to strains and **W** ∈ *M*_*n,r*_(*R*) a cofactor incidence matrix containing covariate information about the strains (e.g., geographical origin, lineage, etc.). Let **x**_*j*_ ∈ *R*^*n*^ be a vector of a genetic variant *j*. The one-component LMM to test the effect of *j*, is typically defined as:

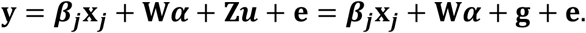

The first two terms are defined as fixed effects, with *β*_*j*_ the effect of the variant *j* and *α* ∈ *R*^*r*^ the vector of effects for the cofactors. The third term **g = Z***u* is a random vector corresponding to the polygenic effect which models the effect of the whole genome (i.e. the genome described by described by **Z**). Because we assume the vector of effects of each genetic variants is the realization of a gaussian distribution *u* ∼ *N*(0_*p*_, *τ***I**_*p*_), it follows that **g** ∼ *N*(**0**_*n*_, *τ***K**) where 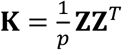 is the genomic relationship matrix GRM. Lastly 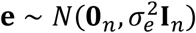 is a random term corresponding to the residual effect. Existing LMM bacterial GWAS typically use the GEMMA software^24^ to estimate the variance component parameters and test the null hypothesis *H*_0_: *β*_*j*_ **=** 0 against the alternative *H*_1_: *β*_*j*_ ≠ 0. An implicit hypothesis of such model is that all strains are extracted from a homogeneous population.

To address heterogeneity in the population studied, the model can be extended to multiple genetic components (MLMM):

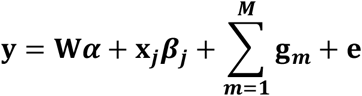

where for all 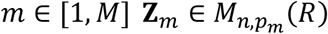 is a (standardized) incidence matrix for the *m*^*th*^ individual random effect, 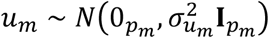 is the the *m*^*th*^ vector of individual random effect, 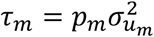 is the variance parameter associated to the *m*^*th*^ polygenic effect and 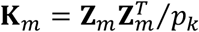 is the *m*^*th*^ kinship matrix. Our *ChoruMM* approach implement such a MLMM, allowing different effect across different groups of strains. Let *c* ∈ ⟦1, *M*⟧^*n*^ a vector partitioning the strains into *M* independent groups and *C*^*m*^ **=** [*i* ∈ ⟦1, *n*⟧ ∣ *c*_*i*_ **=** *m*] the subset of strains classified as *m*. We consider two types of random genetic components: a single shared component capturing genetic variability shared across all strains, and strain-specific components, capturing genetic variability specific to the subset of strains *C*^*m*^. Those components are derived in two steps: first, the unitigs are classified according to their variance within each cluster of strains, resulting in unitigs defined as i) invariant within cluster, ii) variable within exactly 1 cluster, iii) variable within 2 or more clusters. Second, each set of unitigs is used to derive a cluster specific sub-matrix constituting each component of the MLMM. The final multi-component model is expressed as:

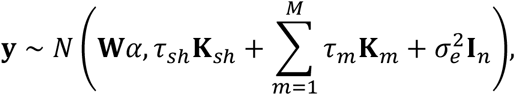

where **K**_*sh*_ is the GRM computed using only the unitigs classified as shared, and **K**_*m*_ are sparse matrices defined as 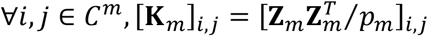 and 0 otherwise, with 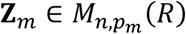 standardized matrices of unitigs containing only the strains in *C*^*m*^ and the unitigs classified as specific to the cluster of strains *m*.

Figure 2 illustrates the proposed pipeline for two and three strain clusters, corresponding to three- and four-components models, respectively: one overall covariance, and either two or three covariances specific to the main branches of the hierarchical plot. Further description of the derivation of the components, and additional details about covariate adjustment, including an algorithm for outlier detection (**Fig. S3**) are provided in the **Methods** section. Finally, for the selection of the number of components, we propose a stepwise approach where an increasing number of components are estimated, and the best model is selected based on the maximum silhouette coefficient. In our Listeria dataset, we observed two maximum that almost perfectly match the lineage and clonal complex (CC) clustering (see **Methods** and **Fig. S4**).

**Figure 2.**
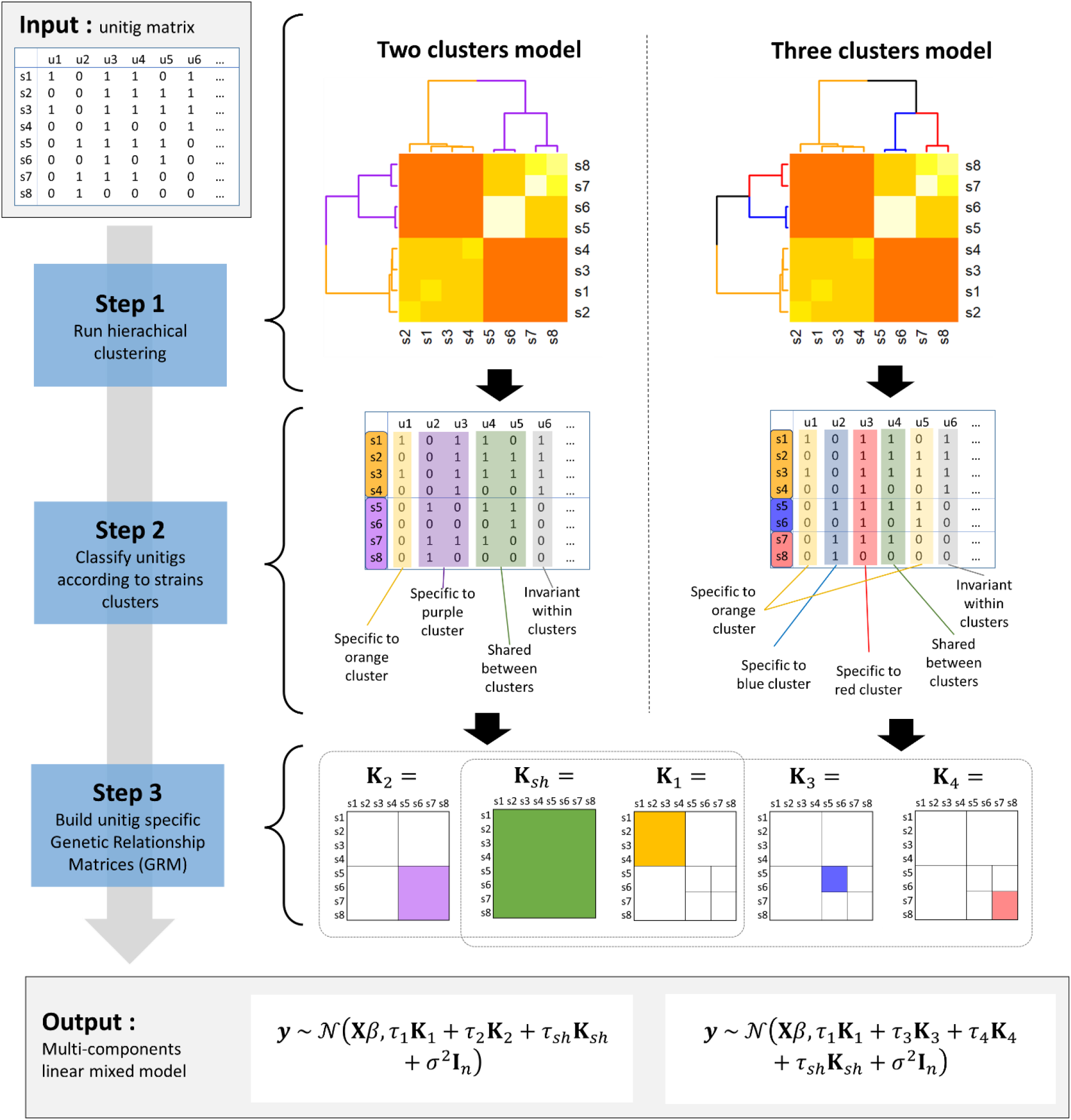
Multi-components LMM pipeline. The input is a matrix of unitigs. In step 1) we compute the genetic relatedness matrix of unitigs and use a hierarchical clustering to form clusters of strains. Unitigs are classified in step 2) according to their variability within strains clusters. In step 3), GRMs are derived for each set of unitigs. The output is a multi-component linear mixed model, with unitigs assigned to a unique genetic term.

### Variance component, type I error and power in simple model

We conducted a series of genetic association analyses using real *Listeria monocytogenes* sequence data and simulated phenotypes to assess the performance of the proposed approach. We first examine the variance component estimates of both our multi-component model and the one-component model when generating either a single overall structure or distinct structures conditional on lineage I and lineage II. Overall, the classic one-component mixed model correctly estimated the variance components for the low heritability scenario but overestimated the genetic component for simulations with strong genetic signal (**Fig. S5**). We argue that this overestimation may be due to the strong structure in the unitig matrix. On the other hand, our multi-component model does not show bias in the variance estimation, whatever the simulated values.

We next estimated the calibration of the type I error by computing the false positive rate (FPR) and the genomic control factor *λ* across four models: i) standard fixed effect not accounting for structure, ii) LMM modeling structure at the cohort level (one component) ; iii) LMM modeling structure at the lineage level (3-components) ; and iv) LMM modeling structure at the clonal complexes (CC) level (16-components). While all tests display a calibrated average FPR, it also exhibits considerable variability across simulations, with a strong deflation of the median and multiple outliers displaying severe inflation (**Fig. 3a**). The *λ* value display similar noisy behaviors in these simulated data (**Fig. S6**). This variability appears to be due to the strong correlation between unitigs (**Fig. 1b**) and disappears when removing the correlation through a permutation of the unitigs (**Fig. S7**). This severely hampers the use of tools commonly used to validate human GWAS results i.e., a visual inspection of the quantile-quantile (QQ) plot and the quantification of test miscalibration through *λ*. Filtering out unitigs based on pairwise squared correlation (*r*^2^) reduces the variability of both the FDR and *λ* value and improves the median (**Fig. 3b** and **Fig S6b**). However, because of the low number of unitig left after filtering (e.g. p = 2061 for *r*^2^ < 0.2, **Fig. 1b**), both FPR and *λ* value estimations display large variance, reducing their practical utility.

**Figure 3.**
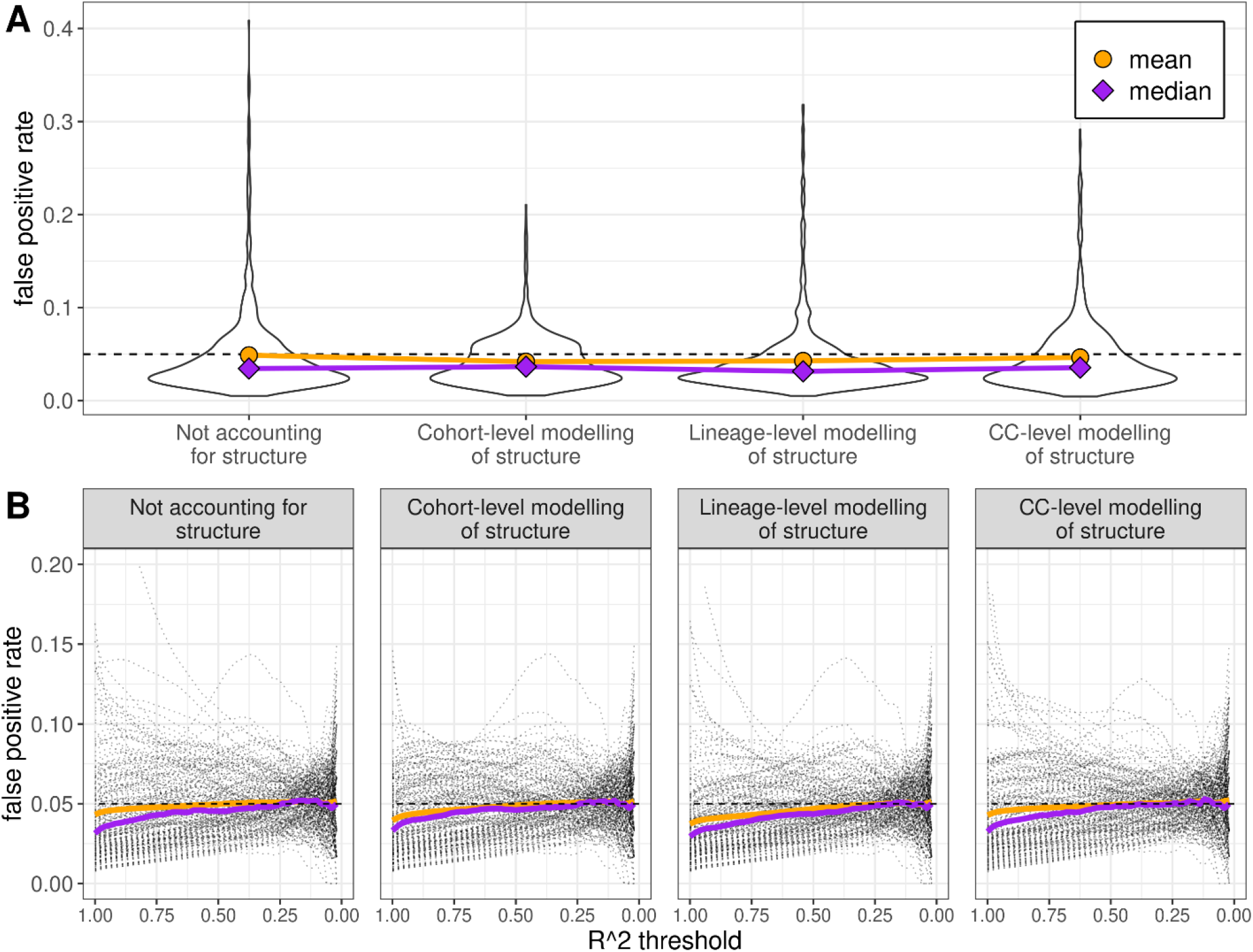
Impact of the strong correlation between unitigs on the p-value distribution. We simulated random phenotypes and for each phenotype we ran GWAS of real unitigs from 907 strains using four linear mixed models (LMM): not accounting for population structure, and accounting for structure at the whole cohort level, the lineage level, and the clonal complex (CC) level. Panel A) showcase the impact of the strong correlation between unitigs on the false positive rate (FPR) over 1,000 simulated phenotypes and 2,000 unitigs. It shows the distributions of FPR plotted using violin plots, with their mean (orange) and median (purple). In panel B) we gradually removed the correlation between unitigs by applying various R^2^ thresholds and derived the FPR (black dotted curves). Also shown are the mean (orange) and median (purple) FPR as a function of the threshold value. For each threshold we simulated 200 phenotypes and tested up to 137,981 unitigs.

Correlation between unitigs can in theory be accounted for through a joint modelling of all unitigs. However, such an approach faces statistical challenges when the sample size is smaller than the number of unitigs and can be computationally prohibitive when the total number of unitig reaches hundreds of thousands, as it requires inverting the unitig correlation matrix. To address these limitations, we propose an efficient empirical solution that, instead of attempting to derive joint estimates, derive the expected distribution of correlated statistics using permutated phenotype values. This null distribution of *p*-values allows to derive 1) a confidence interval for the QQ plots, and 2) a permutation test of the *λ* value to assess deviation from the expected under the null. As illustrated on **Figure S8**, this empirical distribution can be derived using the very computationally efficient fixed effect regression and does not have to be derive using the much more time-consuming mixed models-based test.

Lastly, we conducted a power comparison across LMM tests accounting for structure at either the whole-cohort level, or the lineage level. We simulated series of phenotypes using polygenic models with total heritability *h*^2^ in [0.5; 0.9], and the number of causal unitigs, selected randomly and independently of the structure in the data, in [100; 2000]. As shown in **Figure 4**, despite phenotypes were drawn without assuming lineage or CC structure, we observed a significant increase in power for LMM models accounting for structure at either the lineage level or CC level, as compared to models accounting for structure at the whole-cohort level. For a fixed heritability, the gain in power increases with the polygenicity of the model, when assuming more causal unitigs with smaller effects (N=2,000 vs N=100).

**Figure 4.**
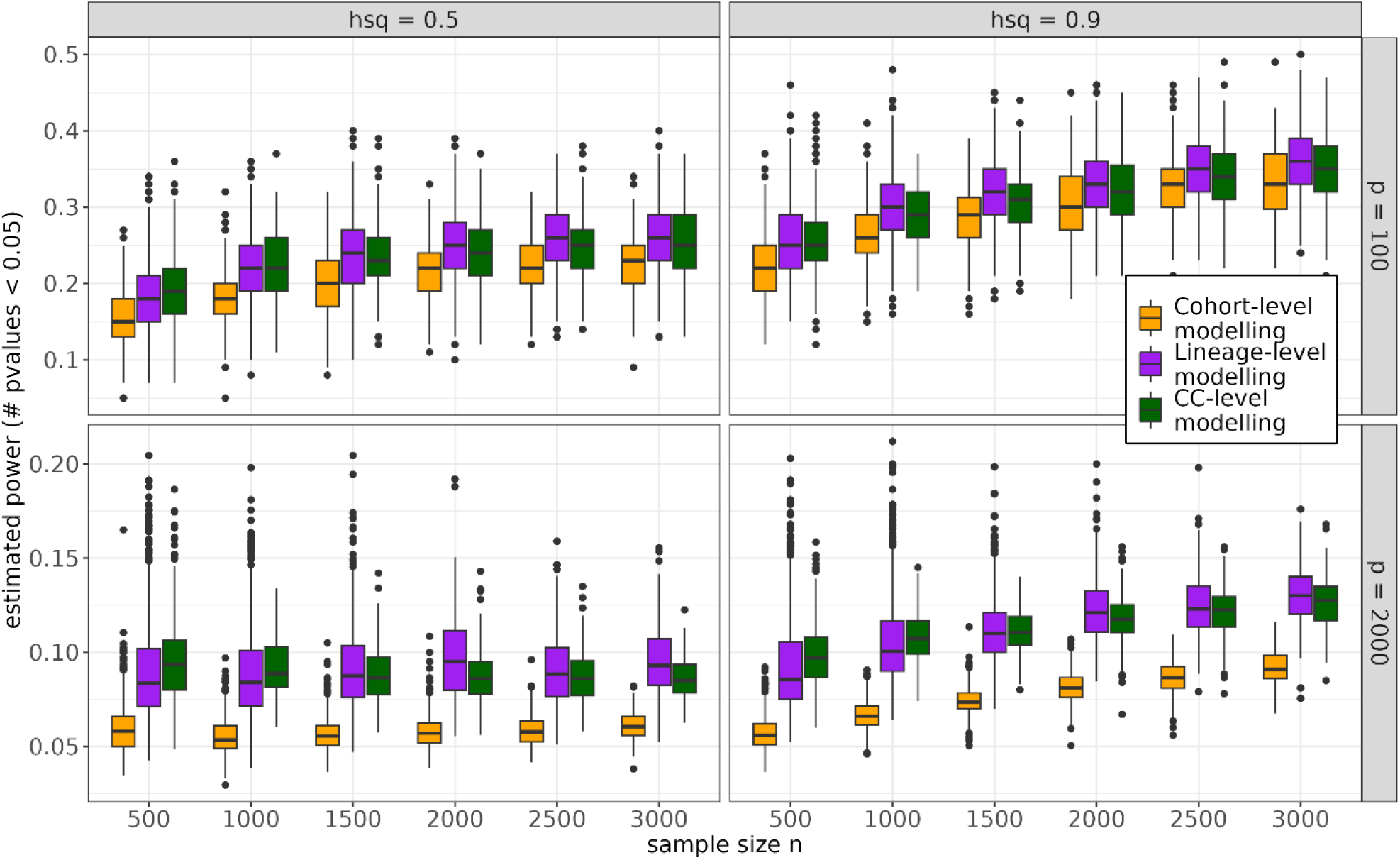
Fine accounting of population structure boosts statistical power. Statistical power was estimated for simulated phenotype with heritability *h*^2^ in [0.5, 0.9] and sample sizes in [500, 1000, 1500, 2000, 2500, 3000]. We first randomly selected subsets of respectively *n* strains and *p* [100, 2000] unitigs, then simulated phenotypes under a linear model with heritability value of *h*^2^. We lastly ran tests for all unitigs, while accounting for the population structure at either the whole-cohort level, the lineage-level or the cohort. Power was then estimated by the mean number of tests with a p-values under 5%.

### Performances in the presence of complex genetic and environmental structure

In the previous section, we considered a fairly simple model, where the genetic effect was drawn independently from the actual genetic structure and assuming the residual noise was homogeneous across strains. To illustrate the potential of our approach, we investigated two additional models and assessed the performances of *ChoruMM* against the state-of-the-art one component model as implemented in *DBGWAS* and *pyseer*, and a simple fixed effect linear regression. First we simulated phenotypic data under the null but in the presence of residual noise confounding the genetic structure. In practice we draw 100 phenotypes as a mixture of standardized normal distribution and the first principal component of a sub-matrix of 22,673 unitigs displaying variability only in lineage II. We tested the association between each phenotype of the 22,673 unitig using the three approaches. The resulting *P*-value distribution over all simulations are presented in **Figure 5a**. As expected, the standard regression is fully confounded, displaying extreme type I error inflation. The one-component model is able to capture part of the confounding effect, but still display substantial inflation, with a miscalibrated *P*-value distribution. *ChoruMM* show the best calibration in this scenario.

**Figure 5.**
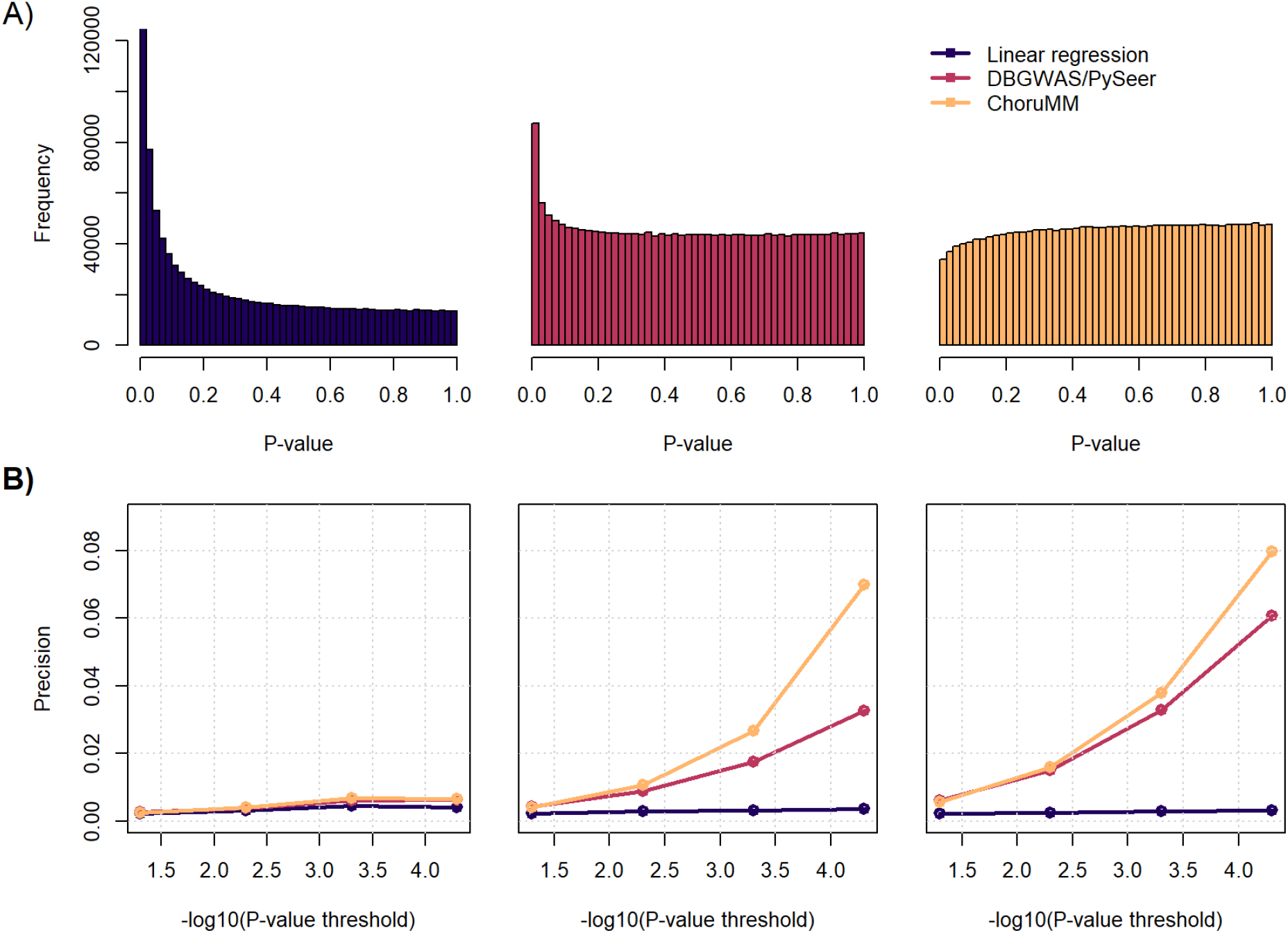
Type I error from real-like data. We compared the performances of three approaches, standard linear regression, one-component LMM as defined in DBGWAS, and *ChoruMM*, in the presence of complex structure under both the null and the alternative. For simplicity, we only model in ChoruMM the lineage effect in a three-component LMM (shared effect, lineage I, and lineage II). A) Histogram of P-value for the three approaches on phenotype simulated under the null but including the effect of confounding factor associated with variability in lineage II. This simulation mimics a scenario commonly assumed in human genetics data. The plot show a an extreme miscalibration of the standard linear regression, a still severe miscalibration for the one-component, and a slight deflation, but no increase in the type I error rate for ChoruMM. B) Precision of the three approaches when testing all available unitigs, in a model where the genetic effect on the phenotype is drawn using 100 unitigs randomly chosen across those displaying variability in Lineage II. Precision is derived as the ratio of true positive signal identified average over the total number of positive signals over 100 simulated phenotypes with heritability *h*^2^ equals 0.1 (left), 0.5 (middle) and 0.9 (right).

We then investigated the ability of our model to decipher true causal unitigs from unitigs associated with the phenotype through correlation with the genetic structure. Here we simulated genetic effect on 100 phenotypes as a function of 100 unitigs randomly chosen across the aforementioned set of 22,673 unitigs displaying variability only in Lineage II. The effects of the causal variants were drawn from a centered normal with variance set so that the heritability *h*^2^ in Lineage II strains equals [0.1; 0.5; 0.9]. We applied the three approaches to all available unitigs mimicking a GWAS analysis. **Figure 5b** presents the average of the precision, defined as the ratio of true positive signal identified over the total number of positive signals, average over the 100 phenotypes. *ChoruMM* show a substantially higher performance with up to two-fold increase in precision when using a stringent significance threshold accounting, typical of GWAS approaches.

### Visualization and statistical significance

Human GWAS results are typically displayed as Manhattan plot that represent the genome-wide - log10(*p*-value) as a function of the chromosomal position. The hallmark of Manhattan plots are peaks of signal around associated regions due to the high correlation between neighboring variants. By construction, unitig have no clear physical position, and a given unitig can potentially display high correlation with any other unitig in the matrix. Consequently, a naïve Manhattan plot for unitigs association results presents disbursed clouds of points, substantially reducing its interpretability. Hierarchical clustering of the unitigs might allow to regroup distant (at the sequence level) but related (at the pattern level) unitigs together, but is computationally prohibitive in large datasets because it has complexity cubic with the number of unitigs. To circumvent this issue, we propose a fast re-ordering algorithm of the unitigs conditional on their pairwise correlation (**Fig. S9**). As illustrated in **Figure 6**, this allows aggregating correlated signals. We propose to further enrich this adapted Manhattan plot by summarizing the distribution of unitig frequencies across CC clusters. Cluster information is multidimensional and complex. As a trade-off between information content and readability, we focused on the mean and variance of the unitig frequencies, which we integrated as a two-dimensional gradient of color. This provides two key features along the association signal: i) the divergence in frequency of each unitig across CC, and ii) the average frequency of each unitig across all CC (**Fig. 6)**. Finally, to account for the extreme correlation between unitigs, we propose an adjusted Bonferroni threshold defined as *α*, the alpha error, over *p*_*eff*_, the effective number of parameters estimated.

**Figure 6.**
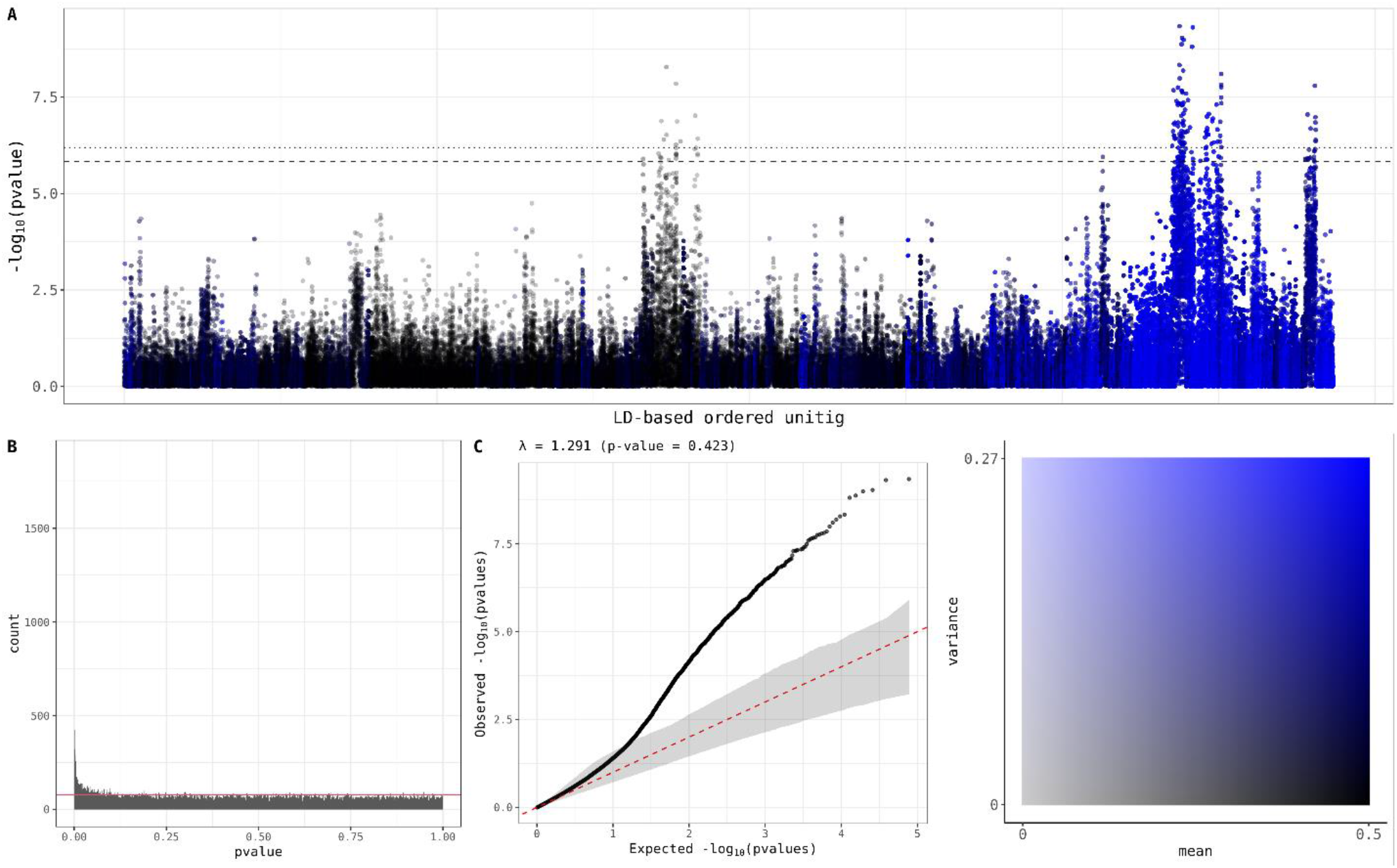
Visualization of GWAS results. An example of GWAS results for a phenotype under the alternative hypothesis. The phenotype was simulated as described in **Fig. 5b** with a heritability of 50%. A) Manhattan plot displaying the -log10(P-values) of each unitig ordered based on the proposed clustering algorithm. The color of each point is computed using both the mean and variance of each unitig frequencies across the strain cluster in the dataset (here 15 CCs) and resumed as a two-dimensional gradient of color. Red points correspond to unitigs with non-null signal. The doted and dashed lines correspond respectively to bonferonni and adjusted bonferonni threshold. B) Histogram of the p-value. C) Q-Q plot of the p-value. The computation of both the confidence interval and the test of *λ* were conducted using 1000 simulations.

## Discussion

Existing bacterial GWAS studies have focused on translating and adapting the association screening aspects of human GWAS but only seldom discussed other features of the genetic modelling. In this study, we propose the use of a multi-component linear mixed model, based on unitigs data, modelling different effects for strains in different biological clusters and accounting for the strong population structure in bacteria. Our model showed control of the type I error once accounting for the strong correlation between variables, while also providing higher power than the classical one-component linear mixed model. Our tool partly addresses the question of deciphering causal variants from passenger mutations because of long-range linkage disequilibrium, an issue pointed early on as a central issue in bacterial GWAS^7,25^. Several reviews also pointed the need to develop visualization tools that, like the Manhattan plot and Q-Q plot from human GWAS, allow to comprehensively explore top association results and quantify possible miscalibration of the test^6,7^. Our study provides a first hint at these challenges. By using a fast algorithm to cluster unitigs, our pipeline provides a Manhattan plot featuring correlation patterns and peaks of signal. We also provide a QQ plot in which the impact of the correlation between variable is accounted for, allowing for visual check of a potential type I error rate inflation. Finally, we tackle multiple testing by controlling for the family-wise error rate using an adapted Bonferroni procedure. All these features have been implemented in the *ChoruMM* package, addressing the need for an end-to-end user friendly unified bacterial GWAS framework^7,25^.

Our approach also has some limitations. First while the *k*-mer covers a broad range of variations, it does not model copy number variation. Note that other recently proposed methods share similarity with our work, but require to analyze separately multiple type of variants (gene presence/absence and SNP within the core genome)^26^. While the approach was developed to be used with unitigs data, it should easily be applied to other genetic variants. Second, the proposed unitig re-ordering algorithm allows to generate a proxy for a Manhattan plot and highlight the main association results. However, the current ordering depends on the data and the selected anchor’s unitig, which makes comparison across studies difficult. A possible path forward would be to define anchor’s reference that can have the same role as core genome assembly. Third, our approach has been developed for continuous outcomes and applicability to binary, while theoretically possible, must be investigated. Fourth, we did not investigate the potential predictive power of the model produced by our approach. For pure prediction purposes, alternative strategies might be considered, including penalized regression models^25^ or machine learning^8^.

## Methods

### The Listeria cohorts

The *Listeria monocytogenes* sequencing data is a non-redundant collection of isolates of food (*n* = 4,551) and clinical (*n* = 2,791) origin collected in France between January 2005 and October 2013. The isolates were collected by the French National Reference Center (NRC) for *Listeria* and the French National Reference Laboratory (NRL) for *Listeria*. Food isolates were collected from multiple sources: 3,143 (69.1%) from food alerts, 178 (3.9%) in the context of investigations following neurological forms of listeriosis, 692 (15.2%) from checks by food industries, and 538 (11.8%) from food surveillance activities. Here we focused on two subsets that passed stringent quality control, in order to limit potential confounding effect. The first, and most stringent dataset, includes 912 Listeria strains that were collected and stored using the same conditions and for which sequencing data were produced using the same protocol. This dataset was used for both the real data analysis and in-depth assessment of our method’s type I error rate. The second dataset, a total of 3,824 Listeria strains. This second dataset is slightly more heterogeneous with variability in the pre-processing of the sample. This dataset was used for power assessment, thanks to its larger sample size.

### Building the unitig matrix

We defined the genetic variants using unitigs built from a De Bruijn Graph (DBG). The DBG offers an exhaustive synthesis of genetic variability across sequences in a cohort of strains. Concatenating nodes within a unique path to obtain the compacted De Bruijn Graph (cDBG) further provides a node incidence matrix of binary variables (with a value of 1 when the node is present in the strain and 0 otherwise) that share strong similarity with the single nucleotide polymorphisms (SNP) matrix of human GWAS. For each bacterial dataset, we derived the DBG using the GATB C++ library^27,28^ and derived the 32-mers from the raw sequence data. The choice of using the 32-mer was driven by recent reviews suggesting it provide an optimized accuracy over computational cost ratio^7^. We start from contigs rather than reads and, consequently, we do not need to filter out low abundance *k*-mers, allowing for the exploration of any variation present in the input genomes. We then compact the DBG using a graph traversal algorithm, which identifies all linear paths in the DBG —each forming a unitig in the cDBG. During this step we also associate each *k*-mer index to its corresponding unitig index in the cDBG. We applied the cDBG pipeline to both *Listeria* datasets, which resulted in a total of 432,226 and 621,244 unitigs, respectively.

### Estimation of genetic relatedness

Previous works have proposed to derive the genetic covariance matrix, commonly referred to as the genetic relationship matrix (GRM), using the same approach as in human GWAS, *i*.*e*. the method of moments proposed by Astle et al ^29^. In brief, the GRM is defined as the product of scaled genetic variants between the pairs of individuals (here the strains) considered, summed over all variants available in the sample. Consider a unitig matrix **Z** of *p* unitigs and *n* strains with element *Z*_*i,k*_ coding the value of a unitig *k* for strain *i*, and denote *f*_*n*_ the proportion of strains including the unitig *k*. The GRM matrix, **K**, of size *n* × *n* can be formed by deriving the genetic covariance between each pair of individuals *i* and *j*:

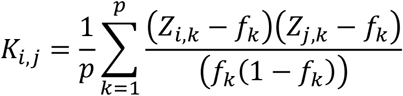

This expression of genetic similarity has been developed on the principle of identity by descent (IBD) for diploid organism, assuming that the relatedness between two diploid individuals can be defined in terms of the probabilities that each subset of their four alleles at an arbitrary locus is identical by descent. It is unclear how a direct translation of this metric to haploid bacterial data impacts the interpretation of genetic similarity. We therefore investigated two alternative derivations of this standard GRM: *K*_*uni*_, a GRM derived using only unitigs present in at least one strain, that is, harboring the combination [(0,1), (1,0), (1,1)] but excluding (0,0) cases ; and *K*_*inter*_, a GRM derived using only unitigs that are shared between the two strains, that is, harboring the combination (1,1). However, while we found some potentially interesting feature with these alternatives, further simulations showed that the three GRMs produce qualitatively similar results when used in a variance component model. For simplicity, we consequently kept the standard GRM in all analyses.

### QC filtering based on the hierarchical clustering

To limit the impact of strain heterogeneity in the whole sample due to outliers, we developed an algorithm to filter out isolated strains based on the hierarchical clustering. We first cluster strains according to their genetic relatedness using a hierarchical clustering algorithm (by default we apply the complete-linkage algorithm, as it is supposed to isolate outliers, on a Euclidean distance-based similarity matrix of the strains). Outliers are identified and removed using the algorithm detailed in **Figure S4**. It consists of a recursive splitting of the hierarchical tree at this root, resulting in a list of up to 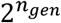 subtrees, where *n*_*gen*_ is the number of times we recursively split the tree. All strains that belong to a subtree that do not contain at least *n*_*leaf*_ strains are considered as outliers in the tree and filtered out. This whole process is repeated until no outliers are left. By default, the hyperparameters of this algorithm, *n*_*gen*_ and *n*_*leaf*_ are set to 3 and 4 respectively.

### Strains clustering and selection of the number of components

The construction of the multi-component linear mixed needs requires to split strain into clusters. The default solution in our pipeline is a data-driven approach based on hierarchical clustering, but the pipeline allows for user-defined clusters (for example lineage or CC). Strains are first clustered according to their genetic relatedness using a hierarchical clustering algorithm (by default we apply the Ward algorithm, as it allows for clean cluster, on the same Euclidean distance-based similarity matrix of the strains as described above). Clusters are obtained by cutting the dendrogram into *K* subtrees of equal height. To pick the optimal number of clusters, we use the average silhouette coefficient, a commonly used metric in clustering that compute how close each strain in one cluster is to strains in the neighboring clusters^30^. It has a range of [-1, 1], with values near +1 and -1 corresponding to respectively well-clustered and potential misclassified strains. For a range of number of clusters, we compute and plot the average of those silhouette over all strains and pick its maximum value as the optimal number of clusters (**Figure S6**). Interestingly two of the partitions of strains which maximize the average silhouette coefficient are composed of either 2 and 15 clusters, which respectively overlap with the observed lineages and CC.

### Unitigs classification

In practice, the unitigs classification is derived as follows. Let *u* ∈ [0,1]^*n*^ denote a unitig and *c* ∈ ⟦1, *K*⟧^*n*^ a vector classifying the strains in *K* clusters. For *k* ∈ ⟦1, *K*⟧ we defined *C*^*k*^ **=** [*i* ∈ ⟦1, *n*⟧ ∣ *c*_*i*_ **=** *k*] the set of strains classified as *k* and:

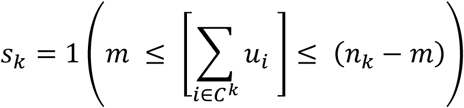

with *m* a hyperparameter corresponding to the minimum unitig count. The *s*_*k*_ should be interpreted as a measure of variability in the cluster *k*; *s*_*k*_ **=** 1 if the unitig is present in *m* strains from cluster *k* and not present in *m* strains, *s*_*k*_ **=** 0 otherwise. If ∑*s*_*k*_ **=** 0, the unitig does not show enough variability within the clustering of strains and is classified as invariant. It will not be included in the computation of any similarity matrix. If ∑*s*_*k*_ **=** 1, the unitig does show enough variability in only one cluster of strains and so is classified as cluster specific. It is included in the computation of the similarity matrix specific to its associated group of strains. Lastly if ∑*s*_*k*_ ≥ 2, the unitig shows enough variability in multiple clusters and is classified as shared. It is included in the computation of the similarity matrix *K*_*sh*_.

### Covariates adjustment and parameters estimation

Additional information from external covariate can be added into the models. Except if specified otherwise, all models include an intercept and two binary variables indicating respectively the lineage (either I or II) and the origin (from food sample or hospitalization) of the strains. CC information was integrated as a random term: denoting by *n*_*CC*_ the number of CCs we first build 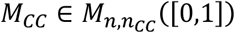 a binarized matrix of the CC, and by assuming a normal distribution of the effects 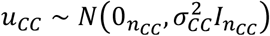, the CC are added to the model as 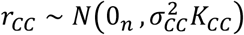 with *K*_*CC*_ computed using the *K*_*all*_ formula on *M*_*CC*_. The estimation of variance components is done by REstricted Maximum Likelihood (REML). Instead of the classic Average-Information REML algorithm, we use the min-max algorithm developed by Laporte et al.^31^, and its implementation in the R package MM4LMM. Significance of unitigs effects is tested using a Wald test, using a *χ*^2^ distribution with 1 degree of freedom under the null.

### Fast re-ordering of unitigs

We developed an algorithm for a fast re-ordering of unitig based on their correlation pattern. A standard hierarchical clustering of the unitigs would be challenging because of a time complexity of *O*(*p*^3^), where p is the number of unitigs. Instead, we propose an approach which assumes that, because of the extreme correlation, the unitig correlation can be captured by random subsets of all available unitigs. Our algorithm has four steps described in **Figure S10**. In step 1, we randomly select *A* **=** 1,000 unitigs referred to as “anchors”. In step 2, the anchors are grouped into non-overlapping sets using hierarchical clustering. As a dissimilarity matrix we first compute 1 minus the squared correlation of the standardized unitig matrix, then compute the *Canberra* distance on this matrix. Hierarchical clustering on the resulting dissimilarity matrix is performed using the complete linkage method. In step 3, we assign all unitigs not in the anchor set to their closest anchor. We achieve this by computing the relatedness between the anchors and the other unitigs, using squared correlation, then assigning every unitig to its most related anchor. Unitigs are ultimately reordered during step 4, in which within-cluster unitigs are sorted according to the R2 with their associated anchors, and clusters are ordered using the dendrogram from step 2. Final unitig order is defined as the concatenation of both within-cluster sorting and cluster ordering.

### Power and type I error simulations

We conducted simulations to study type I error and power for the analysis of continuous outcomes. To assess the type I error, we used a subset of 907 Listeria strains and subsampled sets of unitigs out of the 137,981 distinct unitigs available. Series of random phenotypes were drawn from a normal distribution with mean 0 and variance 1 independently of the unitigs and tested for association using the proposed LMM while considering various models: not accounting for population structure, or accounting for structure at the whole cohort level, the lineage level, or the clonal complex (CC) level. The calibration of the type I error rate was measured using both the False Positive Rate (FPR) and the genomic inflation factor (*λ*), calculated as the ratio of the observed over expected median -log10(*P*-value). In a first series, we drew 1000 replicates, each including a random sets of 2000 unitigs and estimated the distribution, mean, and median for both the FDR and *λ* (**Fig. 3a** and **Fig. S7a**). To assess the impact of the strong correlation between unitigs on those parameters, we conducted a secondary analysis using the same simulated phenotypes but after randomly shuffling the unitigs, in order to remove the correlation (**Fig. S6** and **Fig. S7b**). In a second series, we investigated whether pruning of unitigs *i*.*e*., gradually removing highly correlated unitigs, would reduce the variability in both FDR and *λ*. Here, we drew 200 random phenotypes and tested all unitigs out of 137,981 there were independent at a given R2 threshold that we varied in [1; 0.05] (**Fig. 3b** and **Fig. S7c**).

To assess the statistical power of our approach, we used the extended *Listeria* dataset including 3,481 strains. We randomly sampled 1000, 800, 500, 500, 300 and 200 sets with respective sample sizes in [500; 1000; 1500; 2000; 2500; 3000]. Our algorithm to remove outliers was applied on every subset, and we also filtered unitigs which were not present or absent at least 5 times. For each set we drew phenotypes from unitigs with heritability *h*^2^ in [0.5; 0.9]. Genetic effects were simulated using a standard polygenic model, like those used in human studies. For each replicate, a subset of either *p* **=** [100, 2000] unitigs was selected as causal independently of the CC and lineage status. The effect of each causal unitig *k* was drawn from a centered normal distribution so that *β*_*k*_∼*N*(0, *h*^2^/*p*). The phenotype *Y* was derived as *Y* **=** ∑_*p*_ *β*_*k*_*Z*_*k*_ + *e*, where *Z*_*k*_ is a standardized causal unitig and *e* is a centered normal with variance (1 − *h*^2^). All causal unitigs were tested for association using the proposed LMM while considering various models: accounting for structure at the whole cohort level, the lineage level, or the clonal complex (CC) level. Origin, lineage and CC were also included in the model as fixed effects, when used in variance component.

### Correction for multiple testing

Because of the high number of hypotheses tested in GWAS study, an important number of tested variables will falsely be appeared as associated with the respond. In order to avoid those false positives, one must use multiple testing procedure. The more popular one in human genetic is the Bonferroni procedure, which controls for the family-wise discovery rate (FWER, the probability of having at least on false positive). In practice, it consists of using a stricter significance threshold defined as the level of the test α over the number of tested hypotheses *p*. This procedure supposes all the *p* tested variables are independent, which is not respected as showed previously (and so even then testing only unitig with different pattern). Consequently, a straight application of the Bonferroni procedure will produce a very stringent significance threshold, and potentially interesting signals will be discarded.

To circumvent this issue, others have proposed to rather control for the false discovery rate^19^, which may improve the number of detected signals at the cost of a higher number of false positive. Because discoveries need to be verified via wet experiments, false positive results may lead to potential waste of both time and money and so must be avoid as much as possible. Therefore, we preconize the use of Bonferroni procedure to keep strong control of false positive number through the FWER rate as done in human GWAS. To account for the strong correlation between the tested unitigs and in order to improve discovery potential, others have proposed to use permutation based approach^32^. However, permutation might be computationally expensive, and accounting for the structure in the permutation might be challenging.

Here, we propose the use of an adapted Bonferroni procedure by using the effective number of tests as described in Gao et al^33^ instead of the overall number of tests *p*. Let **M** ∈ *M*_*n,p*_(*R*) an incidence matrix of highly correlated unitig and **Z** ∈ *M*_*n,p*_(*R*) its column-standardized version. The approach consists in computing the associated correlation matrix ***Z***^***T***^**Z**, applying a principal component analysis on this matrix. The effective number of tests can then be estimated as the count the number of principal components needed to reach 99.5% of variance explained. In practice, we split **M** into *l* correlation blocks **M**_*i***=**1…*l*_ derived from our clustering of unitigs and compute the derivation for each block to obtain *p*_*eff*.*i*_. The overall effective sample size is derived as *p*_*eff*_ **=** ∑_*l*_ *p*_*eff*.*i*_

