## Supplementary figure for "*ChoruMM*: a versatile multi-components mixed model for bacterial-GWAS"

### Supplementary material

***Figure S1. Unitig inference***

Illustration of the unitig inference from raw sequencing data for three strains *s1*, *s2* and *s3*. Raw sequence data (a) are first used to build *k*-mers and mapped together using De Bruijn graphs based on overlapping *k*-mers (b). The graphs are then compacted into unitigs (c) and form paths (d). The resulting paths can then be used to build a unitig matrix where presence for a gicen strain is coded as 1 and its absence as 0 (e).

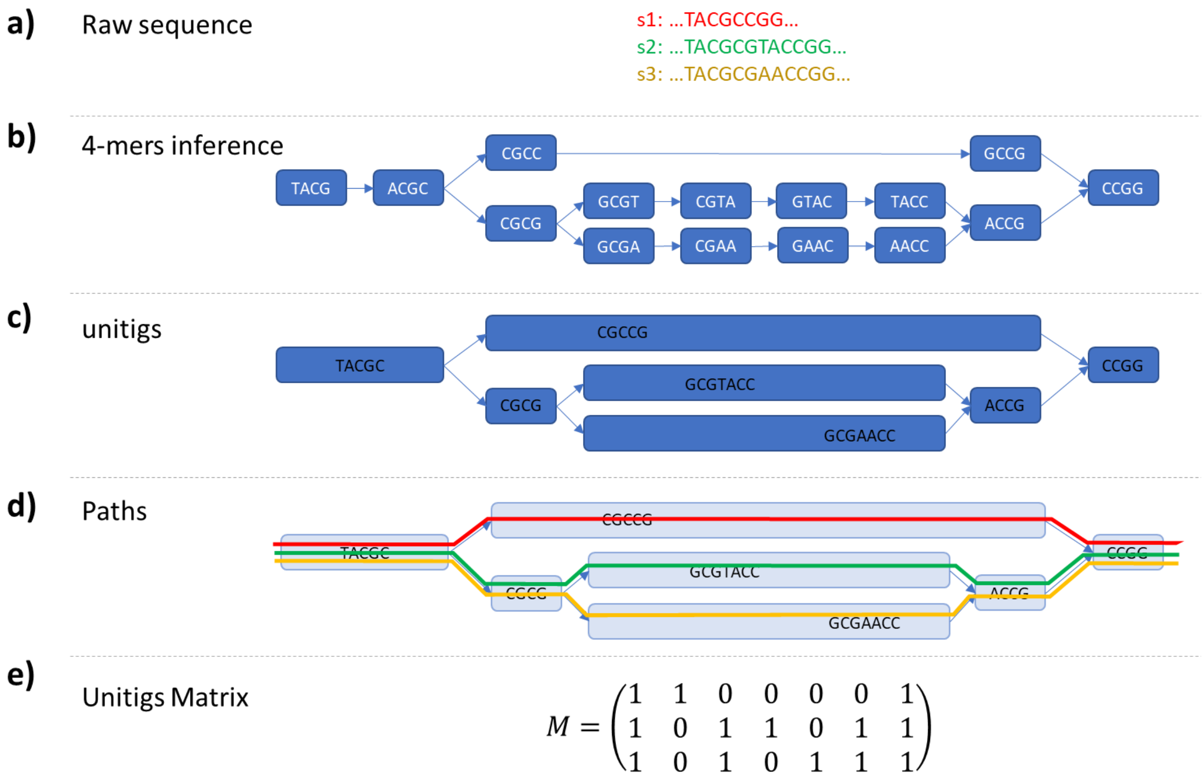

***Figure S2. Genetic variance in lineage I and II conditional across various generative models***

Phenotypes were simulated as a function of sets of 2,000 real unitigs and derived the realized genetic variance in the whole sample (dark blue), in Lineage I (purple) and Lineage II (orange) separately. Unitigs were drawn conditional on the differences in unitig frequency (UF) between Lineage I and Lineage II using three approaches: i) unitigs with a frequency larger than a threshold in the total sample (panel a and d); ii) unitigs with a frequency larger than a threshold in both lineages (panel b and e); and ii) unitigs which difference in frequency between the two lineages is smaller than a given threshold (panel c and f). The true total genetic variance equals 0.5 (top panels) or 0.8 (bottom panels).

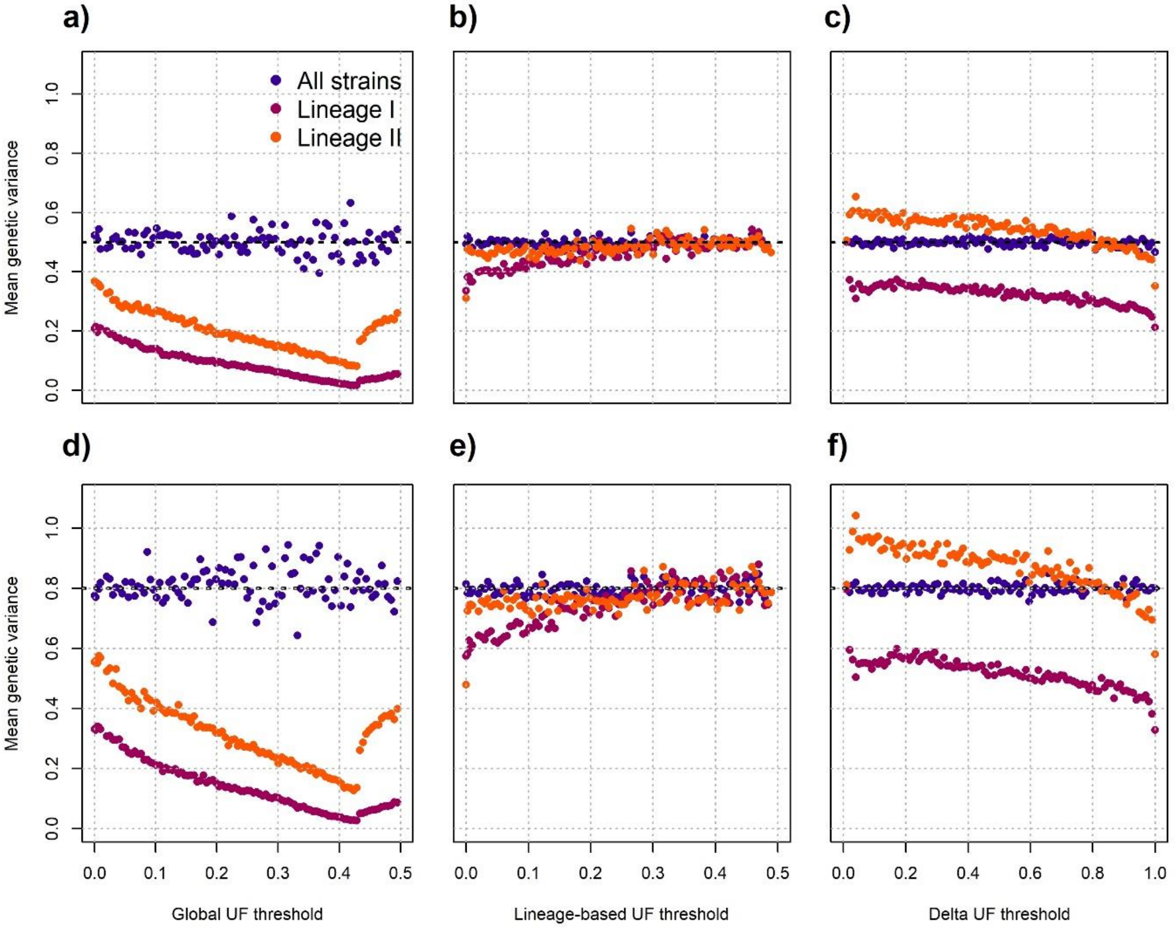

##### *Figure S3. Algorithm to filter isolated strains using hierarchical clustering*

The algorithm we use to remove isolated strains in the dendrogram. Its input is a list of strains and its associated matrix of unitigs. First, we run hierarchical clustering to obtain a tree, which is next recursively splatted at its root. We then compute the number of leaves in every associated subtree, and lasty remove all strains in subtrees without enough leaves. The list of strains in then updated, and the algorithm goes until no strains are detected as outliers.

**input** : $\mathcal{S}$ a list of strains, $\mathbf{U}$ its associated unitig matrix.
**output:** $\mathcal{S}$ updated without outliers.

$n_{out} \leftarrow 1$;
**while** $n_{out} \neq 0$ **do**
    *Run hierarchical clustering for all strains in* $\mathcal{S}$ ;
    $T \leftarrow$ tree from hierachical clustering using $\mathbf{U}[\mathcal{S}]$ ;
    $\mathcal{T} \leftarrow \{T\}$ ;

    *Recursively split* $n_{gen}$ *times the elements of* $\mathcal{T}$ *at their root* ;
    **for** $i \in [\![1, n_{gen}]\!]$ **do**
        **foreach** $T_i \in \mathcal{T}$ **do**
            **if** $\#T_i \neq 1$ **then** $T_i \leftarrow \{T_i^{right}, T_i^{left}\}$ ;
        **end**
    **end**

    *Check if every subtree in* $\mathcal{T}$ *has at least* $n_{leafs}$ *leafs* ;
    **if** $\exists\ T_i \in \mathcal{T} \setminus \#T_i < n_{leaf}$ **then**
        $\mathring{T} \leftarrow \{T_i \in \mathcal{T} \setminus \#T_i < n_{leaf}\}$ ;
    **else**
        $\mathring{T} \leftarrow \emptyset$;
    **end**

    *Remove outliers (if any)*;
    **if** $\#\mathring{T} \neq 0$ **then**
        $\mathcal{S} \leftarrow \mathcal{S}\ /\ \{s_i \in T_i\ ;\ T_i \in \mathring{T}\}$
    **end**
    $n_{out} \leftarrow \#\mathring{T}$
**end**

***Figure S4 Average silhouette coefficient plot***

Average silhouette coefficient against various cluster number derived from real *Listeria* data, derived using the Euclidian distance between the strains then performing hierarchical clustering using the Ward algorithm.

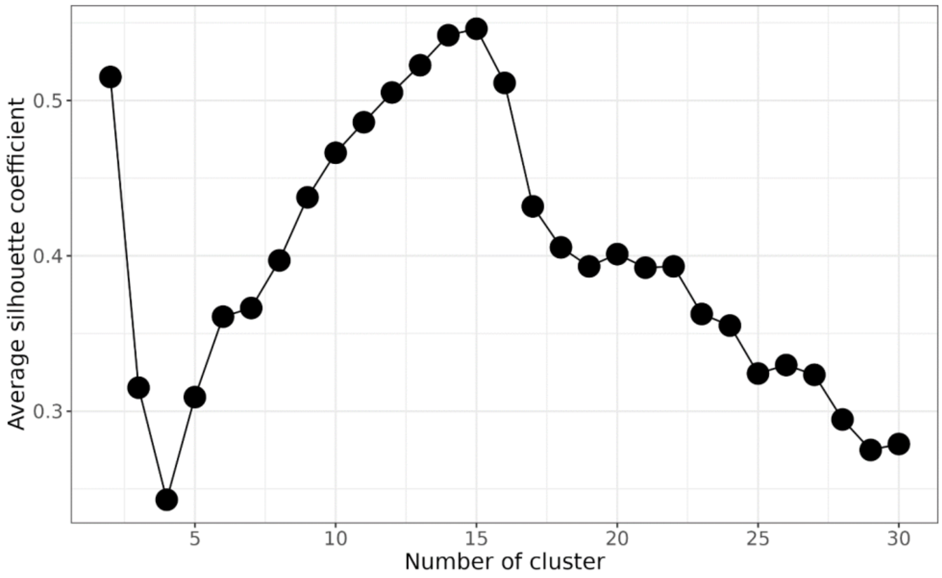

***Figure S5. Estimation of variance components***

Boxplot of estimated variance components. We generated phenotype according to various genetic model (one model per panel, with left and right rows corresponding to model which respectively account or do not account for the strains lineage). Variance components were then estimated using our developed linear mixed model. Every boxplot included 200 simulated phenotypes.

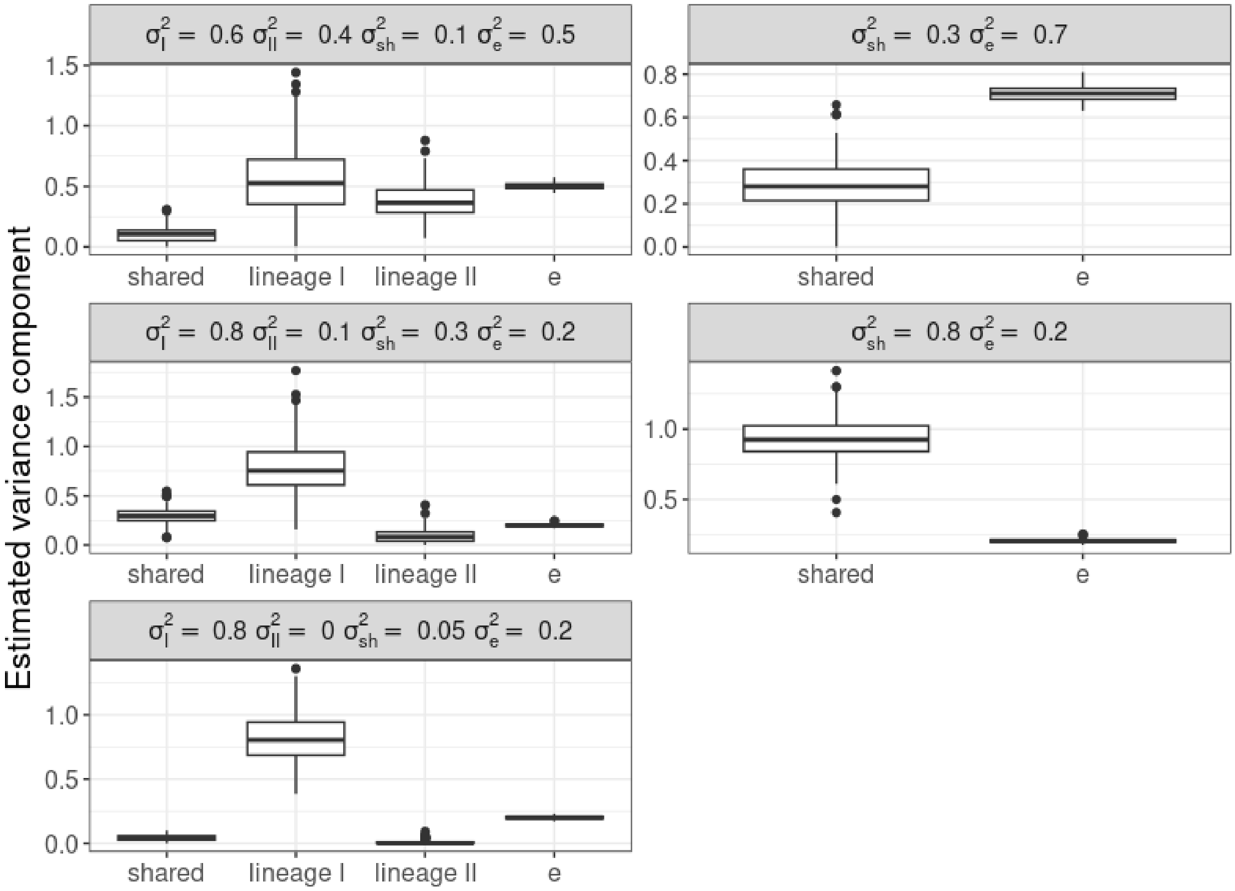

***Figure S6. Impact of unitigs correlation on the genomic inflation factor***

We derived the inflation factor $\lambda$ from the data simulated in Figure 3 (Panels A and C), that uses real unitigs data and unitigs filtered based on their pairwise correlation, and from Figure S6 (Panels B), that uses randomly permuted unitigs in order to break the correlation. Panel A) display the distributions of $\lambda$ along their mean (orange) and median (purple) for the four LMM models considered: not accounting for the structure, or accounting for structure at the whole cohort level, the lineage level and the clonal complex (CC) level. Panel B) display the estimated $\lambda$ when gradually removing the correlation between unitigs by applying an increasingly stringent R-squared threshold. The black dotted curves correspond to the $\lambda$ of one simulation. The orange and purple curves correspond the mean and median $\lambda$ for a given R-squared threshold, respectively.

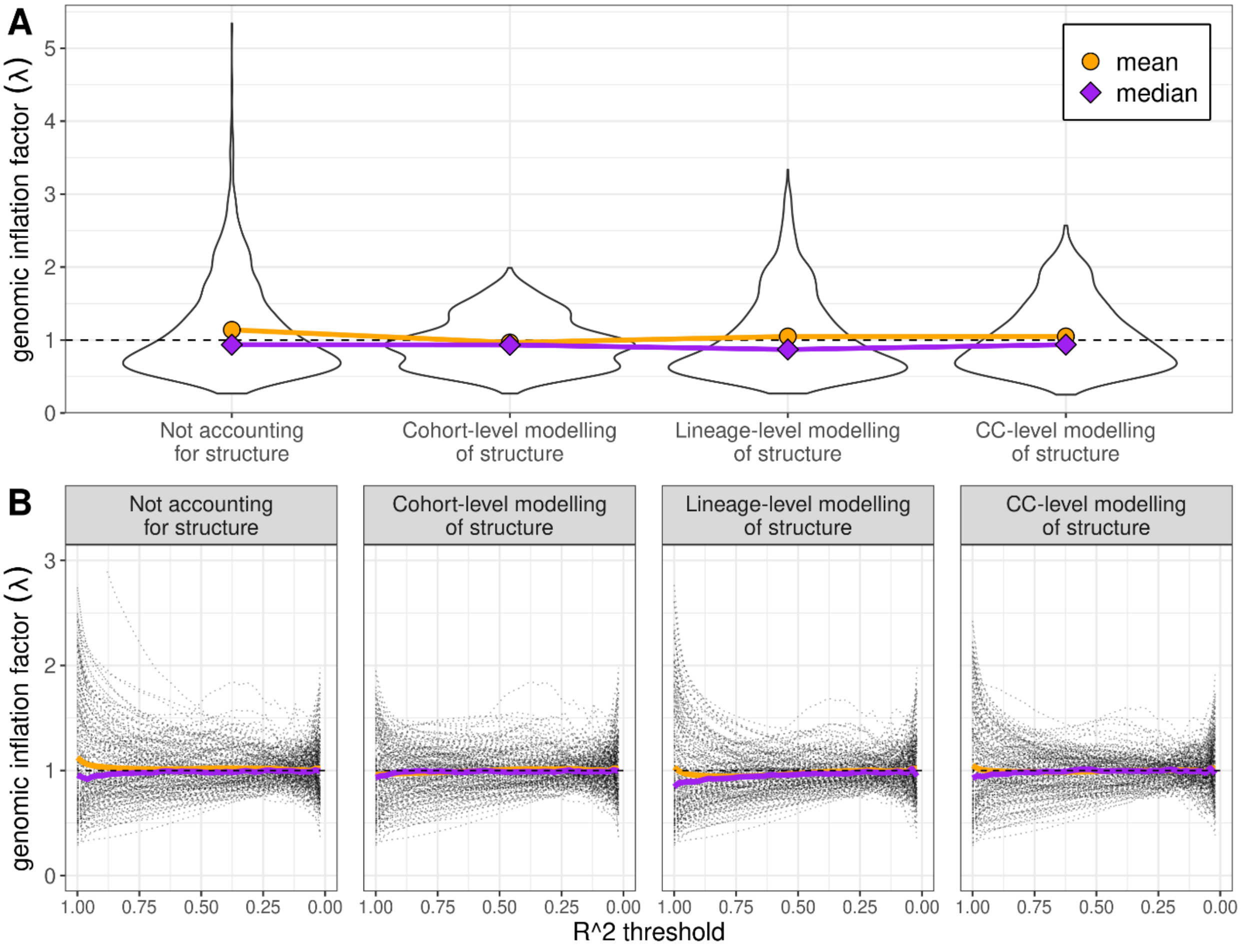

***Figure S7. Type I error rate and genomic inflation factor for non-correlated unitig***

This figure describes the calibration of the type I error without the correlation between unitigs. We simulated random phenotypes, ran GWAS on a subset of 2 000 unitigs with permuted labels (in order to break population structure) for various LMM and computed the false positive rate (FPR) to estimate the type I error. This process was replicated 1000 times. The distributions of respectively FPR and $\lambda$ were plotted in panel A and B, using ridges plots. We also displayed the distributions of correlated FPR and $\lambda$ for comparison. We notice that the FPR is slightly over the test's level when accounting for the population structure, with the bias increasing with the number of random components.

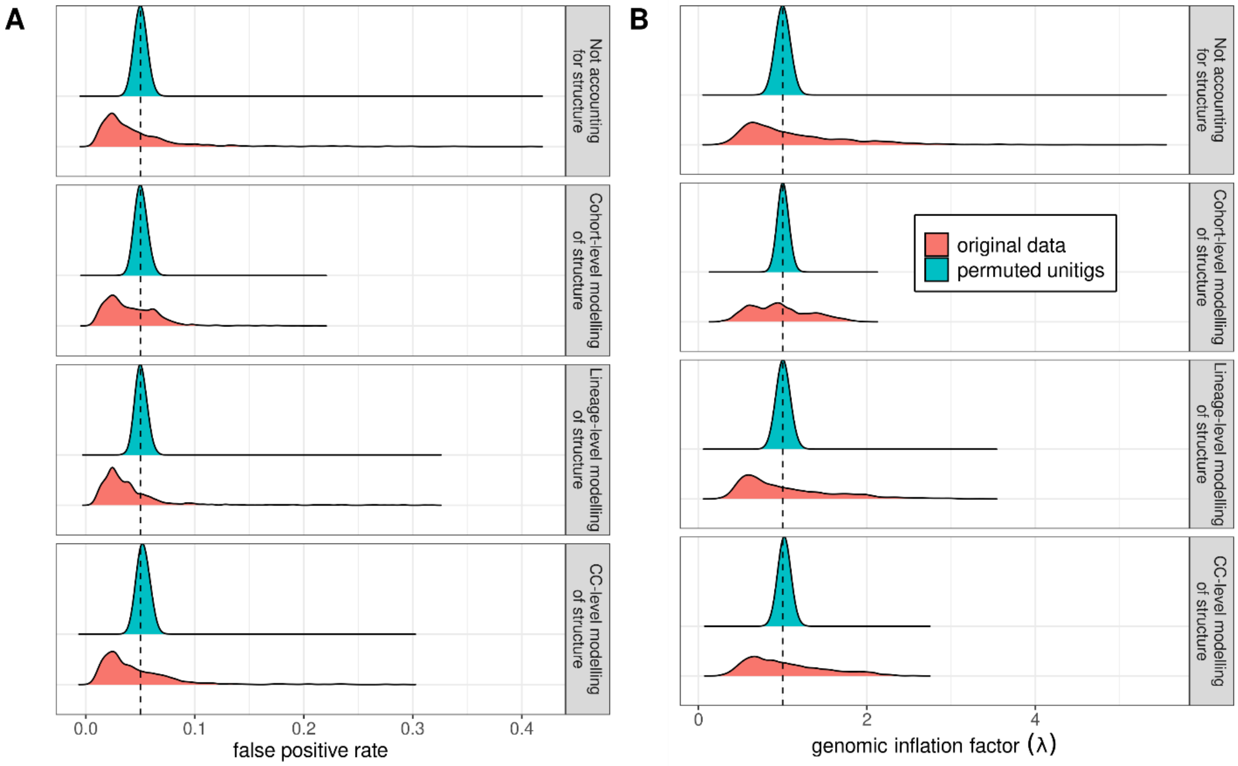

***Figure S8 Empirical QQ plot confidence interval in simulated data***

Panel A represents the QQ plots of the distribution obtain from the simulations used in **Fig 3**. Each panel represent the model used for the testing (Cohort-level, Lineage-level and CC-level mixed model, or no mixed model). Each dotted grey line represents the distribution of one simulation, while the full blue lines correspond to respectively the 2.5% and the 97.5% quantiles. The red full line is the diagonal. B) represents the 2.5% and the 97.5% quantiles obtain from methods accounting for the structure against the respective quantiles obtain when not accounting for the structure. The black dotted line is the diagonal.

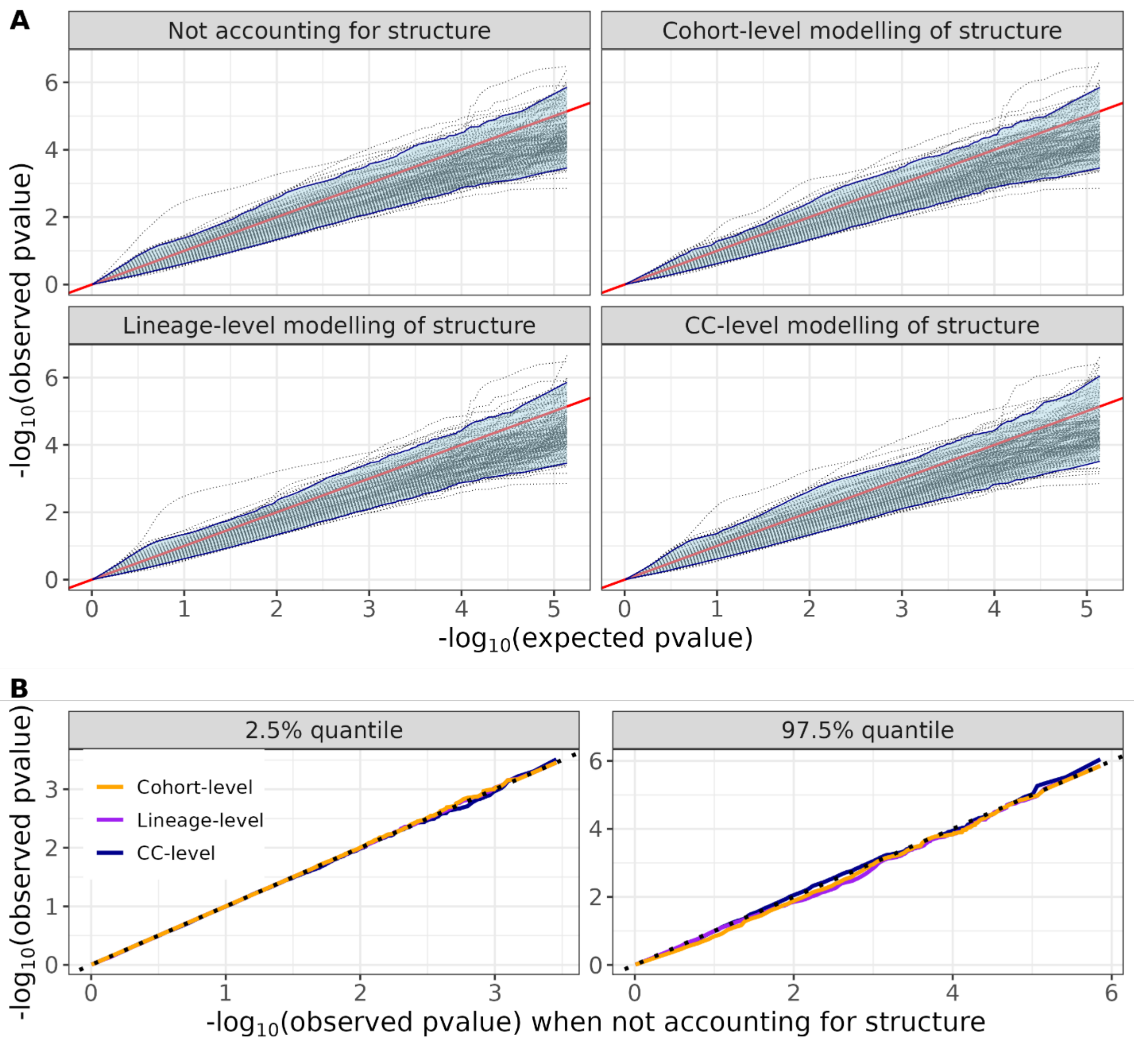

*Figure S9. Fast re-ordering of unitigs*

Pipeline for a fast re-ordering of unitigs based on correlation pattern. A subset of unitigs is randomly selected as anchors in step 1, then clustered together in step 2. Relatedness between the anchors and all the unitigs is computed during step 3, and unitigs are assigned to a cluster according to their closest anchor. Lastly unitigs are re-ordered according to both their relatedness with their associated anchor and the clustering of the anchors.

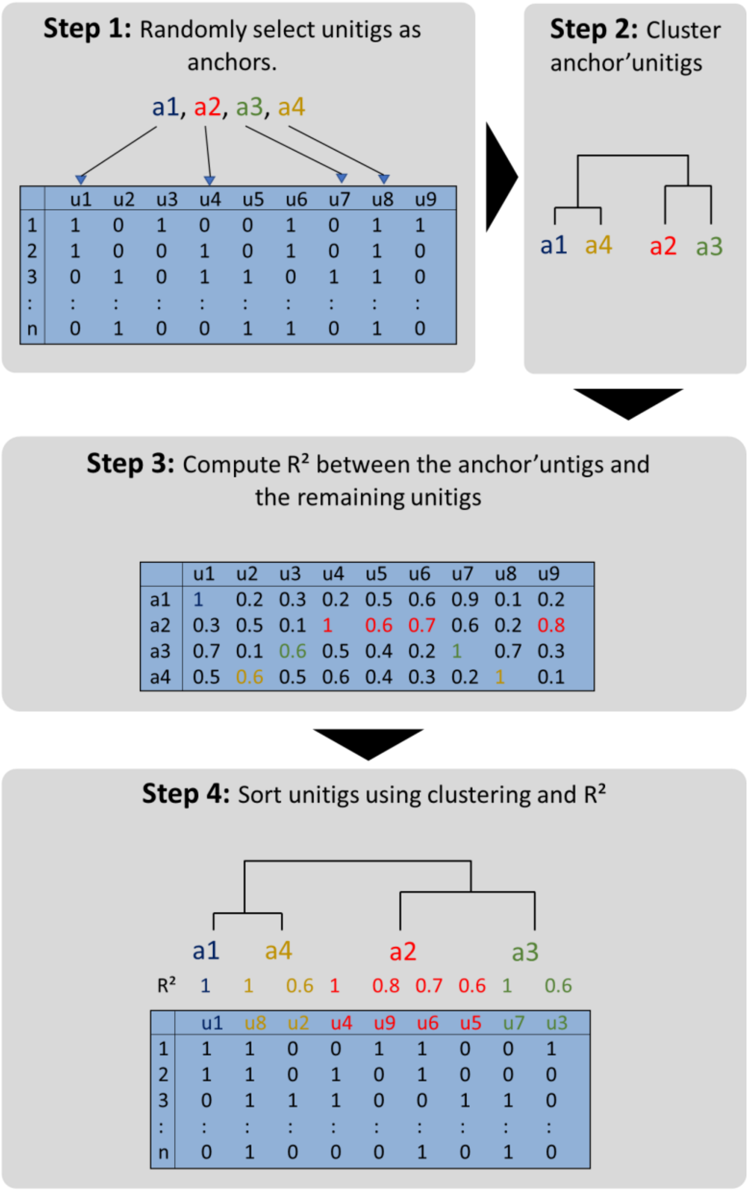
